# Spatiotemporal Divergence Between Intrinsic And Evoked Cortical Activity Predicts Visual Detection

**DOI:** 10.64898/2026.01.06.697966

**Authors:** Dylan Jensen, Zachary W. Davis

## Abstract

The threshold for sensory detection varies with fluctuations in intrinsic cortical activity. When stimulus-encoding cortical populations are more excitable, stimuli elicit stronger neural responses that are more likely to be detected. However, the detection of a stimulus is also more likely when cortical populations are less excitable because there is less background “noise”. Therefore, it is unclear how the variable states of intrinsic and sensory-evoked cortical activity and their interactions impact sensory detection. We hypothesize the answer depends on the spatiotemporal structure of intrinsic activity states across sensory encoding and non-encoding populations. To test this, we examined intrinsic and target-evoked population activity across cortical Area MT in common marmosets while they performed a threshold visual detection task. We compared detection performance based on target-evoked responses and the state of intrinsic activity in the larger surrounding population. We find that the intrinsic activity in the surrounding, non-encoding population predicted trial-by-trial detection performance better than the population encoding the target-evoked response. Furthermore, we find that the detection performance of the monkey was best predicted by the divergence in excitability between the encoding and surrounding non-encoding population. These findings suggest that, rather than a source of noise or irrelevant to sensory processing, the distributed spatiotemporal state of intrinsic activity directly influences how sensory signals are represented in cortical populations and can influence perceptual thresholds in visual detection.

**Significance Statement:** Prior research into how variability in neural activity impacts perception has often focused on neural populations that encode relevant sensory information. However, the role of variable intrinsic activity in nearby, non-encoding populations and their contribution to sensory representations is less well understood. We found that the state of intrinsic activity in non-encoding populations was a better predictor of performance on a threshold visual detection task than the evoked-response magnitude. These results suggest that the state activity in broader neural populations plays a larger role in sensory computations relevant to perceptual decisions than previously regarded.

## Introduction

There is a long tradition in neuroscience examining how variability in sensory-evoked responses among neurons whose receptive fields are tuned for task-relevant features (e.g. location, orientation) are predictive of the perceptual choices made by behaving animals (Celebrini and Newsome, 1994; Britten et al., 1996; Shadlen et al., 1996; Polonsky et al., 2000; Roelfsema and Spekreijse, 2001; Palmer et al., 2007; Chen et al., 2013). In the visual system, identical presentations of a faint or ambiguous visual stimulus produces variable trial-to-trial responses (Tomko and Crapper, 1974; Vogels et al., 1989; Shadlen and Newsome, 1998). The magnitude of the stimulus-evoked responses in neurons tuned for a location or stimulus feature are correlated with the choices made during perceptual decision making tasks (Celebrini and Newsome, 1994; Britten et al., 1996; Shadlen et al., 1996; Polonsky et al., 2000; Roelfsema and Spekreijse, 2001; Palmer et al., 2007; Chen et al., 2013; Crapse and Basso, 2015). However, sensory responses can have limited predictive power for behavioral outcomes (Crapse and Basso, 2015). The poor predictive power of stimulus-responsive neurons has often been attributed to variability in cortical activity across stimulus presentations. Identical sensory inputs elicit stronger or weaker responses contingent on the state of variable cortical activity (Fellous et al., 2004; Schölvinck et al., 2015). Depending on how this variability is correlated across the population along dimensions that define the sensory signal, these variable states can impact the performance of downstream decoders that govern behavioral decisions (Zohary et al., 1994; Nienborg et al., 2012; Moreno-Bote et al., 2014; Nandy et al., 2020).

Consider the problem of threshold visual detection (Fig. 1*a*). The brain must identify the presence of an unpredictable weak visual signal in an environment of both ongoing sensory driven and intrinsic cortical activity. The detectability of the target can be quantified by the separation between the target-absent population activity and the target-present population activity (Ress and Heeger, 2003). Accordingly, stronger target-evoked responses are more reliably detected than weaker target-evoked responses, leading to the idea that visual detection requires activity above some threshold (van Vugt et al., 2018). This threshold might be derived from the state of activity in the broader population because there is evidence that relatively quiet activity states have lower decision thresholds (Parker and Newsome, 1998; Iemi et al., 2017). This creates an apparent conflict: Sensitivity appears to be higher during states when the cortical population that is responding to the target is more excited because that yields stronger evoked responses. But detection performance may also benefit during states when the cortical population is less excited because this reduces the background activity that would otherwise mask weak sensory signals (Wu et al., 2024). How do these variable states of intrinsic cortical activity impact detection sensitivity?

**Figure 1.**
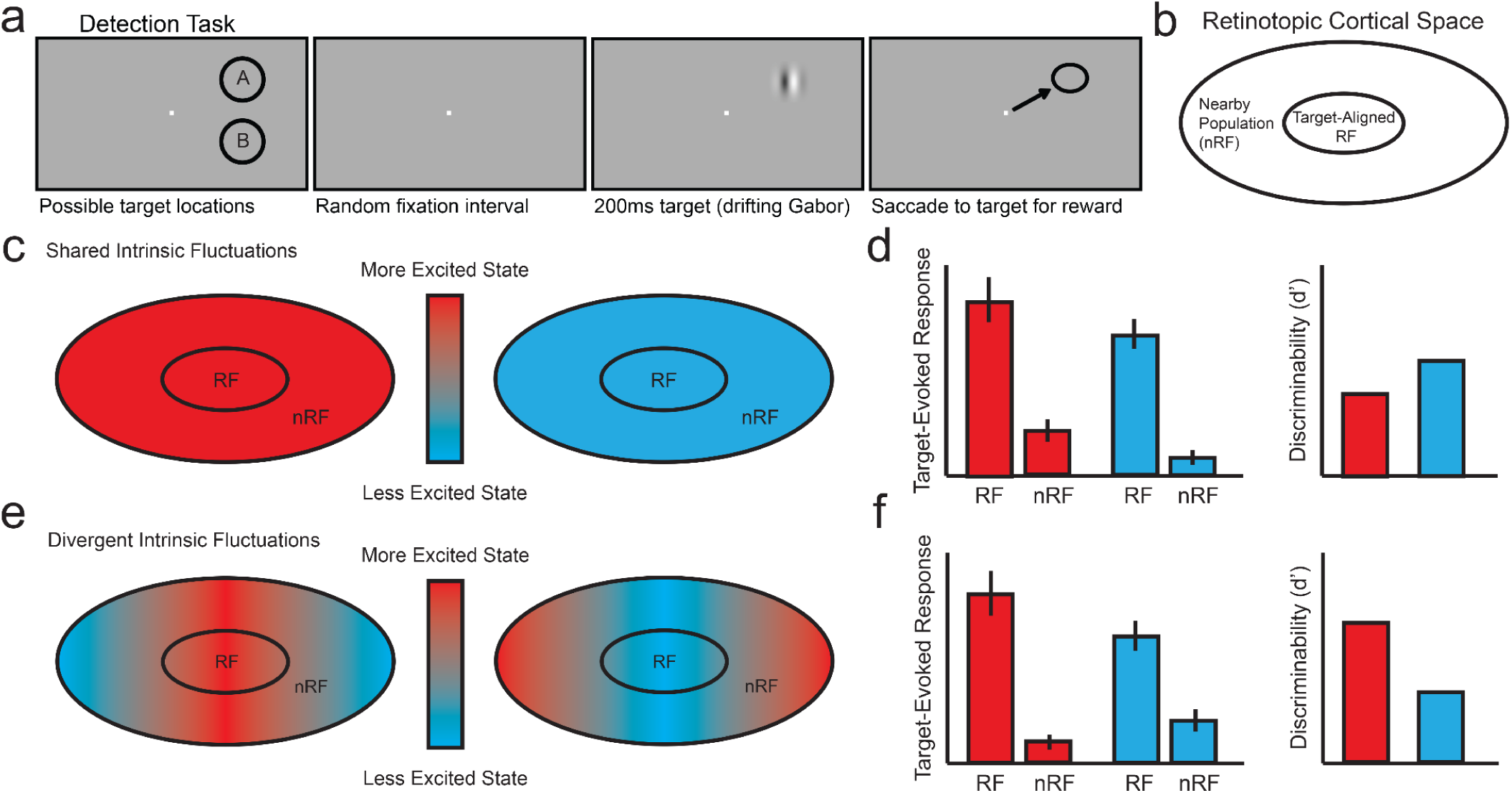
The effect of intrinsic fluctuations of stimulus detectability depends on their spatiotemporal structure. **(a).** Schematic of the visual detection task performed by 2 marmosets. A target is equally likely to appear at one of two target locations at an unpredictable time. Detection of the target is reported with a saccade to the target location for a reward. **(b)** We considered two spatially defined cortical populations in area MT: the neural population retinotopically aligned with the target location (RF) and the nearby population that is adjacent to the RF population but does not exhibit a target-evoked response (nRF). **(c).** Illustration of the shared fluctuation hypothesis. Intrinsic fluctuations in cortical excitability can be shared across the RF and nRF populations as they vary from states of high excitability (red) to states of low excitability (blue). **(d)**. Hypothesis for the effect of shared excitability states on the RF and nRF responses to the target and the discriminability of the target-evoked activity from the ongoing activity of the surrounding population. More excitable states will increase both the RF evoked response and the spontaneous nRF population activity as compared to less excitable states. If the increase is additive and the variance scales with the mean activity, the increase in nRF activity will reduce the discriminability of the RF target-evoked response from the surrounding nRF activity. **(e)**. The alternative hypothesis of divergent excitability states on detection performance: the excitability state varies spatially across the RF and nRF populations. **(f).** The prediction is that excitability states that foster divergence between RF and nRF activity will improve discriminability and opposite states will impair discriminability.

To address this question, we considered two hypotheses regarding the possible differential contribution of intrinsic activity on visual detection sensitivity (Fig. 1*c*, *e*). The first hypothesis is if fluctuations in intrinsic cortical excitability are largely shared amongst both the population whose receptive fields are aligned with the visual target (hereafter called the RF population) and nearby neurons who do not respond to the appearance of the target (hereafter called the nRF population) than an increase in intrinsic excitability will both increase the magnitude of the RF evoked response and the surrounding nRF activity (Fig. 1*d*). This shared increase in RF and nRF activity could reduce discriminability between target-evoked responses and the surrounding background activity resulting in poorer detection performance. Conversely, because the variance in population activity scales with the mean, if the shared intrinsic activity is weaker, then while the target-evoked response may be smaller, the reduction in background activity would reduce the threshold for detection resulting in better performance. Under these conditions, we would predict that detectability is greater when overall intrinsic activity is weaker.

Alternatively, if the states of intrinsic activity in the RF and nRF populations are not shared, but rather exhibit some heterogeneity, then there could be divergence between the intrinsic activity of the RF and nRF populations. When there is stronger intrinsic activity in the RF population and weaker intrinsic excitability in the nRF populations, evoked responses will be stronger and stand out more from a quiescent background environment, resulting in better detection performance (Fig. 1*f*). Conversely, when RF population activity is weaker and nRF population activity stronger, weaker evoked responses will be masked by stronger activity in the background environment resulting in poorer detection performance. While under both hypotheses the best performance occurs when the nRF population is more quiescent, our prediction is that detection performance is enhanced when RF and nRF activity diverges across the two populations favoring the RF neurons. This framework implies that detection performance doesn’t depend solely on the magnitude of the RF evoked response, but rather the interaction between RF and nRF activity at each moment in time.

To test this prediction, we measured the distributed ongoing and visually-evoked activity simultaneously recorded from Utah multielectrode arrays chronically implanted within Area MT of common marmosets *(Callithrix jacchus)* while they performed a threshold visual detection task. We compared the activity of RF and adjacent nRF populations, defined by their encoding of target information, in periods before and during the target-evoked response. We found that detection performance was correlated with the activity difference between the RF population and the nRF population during the evoked response period. Hits were characterized by stronger RF evoked responses in the presence of reduced nRF activity and vice versa for misses. Further, we find that the level of nRF activity and the divergence between the activity of the RF and nRF populations are better predictors of trial outcome compared to the evoked response magnitude in the RF population alone. Finally, we find that this divergence between RF and nRF population activity is not a consequence of the stimulus evoked response, but rather is present in the ongoing intrinsic activity, as indexed by the local field potential (LFP), at the onset of the evoked response. These results highlight the importance of considering the state of the larger population of neurons on behavior as they provide context to the activity of neurons that are tuned for stimulus features in behavioral tasks.

## Results

We analyzed data from two marmosets that were trained to perform a threshold visual detection task (Fig. 1*a*). In the task the monkey had to detect targets that appeared at unpredictable times at two equally eccentric locations in the visual hemifield contralateral to a multi-electrode array chronically implanted in marmoset Area MT (Fig. 2*a*). RF and nRF populations were defined based on the response to each target position in the retinotopic map of the cortical area, which corresponded to two distinct locations on the multielectrode array (Fig. 2*b*, *c*). RF channels were those that showed significant increases in activity with respect to the baseline firing rate in response to the target and nRF channels were defined as those adjacent to RF channels that showed no significant increase in activity (Fig. 2*d, e*). Figure 2*f* shows the average evoked response across the RF and nRF channels as a function of time. There was no significant change in nRF activity during the evoked response period (50-250 msec after target onset) compared to the period just before the target appeared (−300 to −100 msec before target onset; mean normalized firing rate pre 0.96 ± 0.027 v. post 0.98 ± 0.028 S.E.M.; p = 0.65 Wilcoxon signed-rank test; N = 45 sessions).

**Figure 2.**
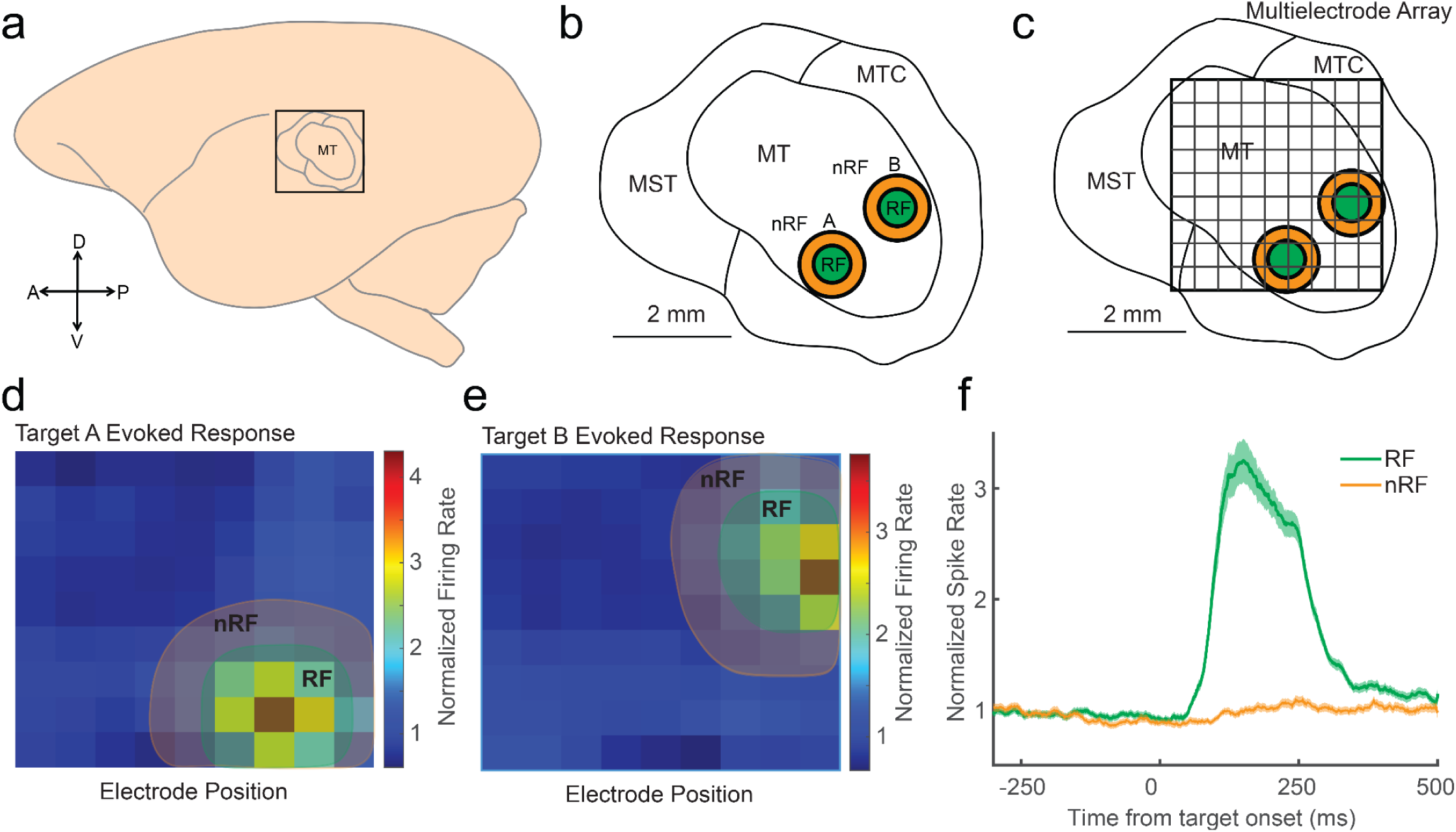
RF and nRF populations defined based on the evoked responses to targets in the detection task. **(a).** Multielectrode arrays were implanted in Area MT of 2 marmosets trained to perform the visual detection task. **(b).** Illustration of Area MT and the positions of the 2 targets (A and B) in the retinotopic map of the cortical area. The RF (green) and nRF (orange) regions for each target location are shown. **(c).** Schematic of the alignment of the Utah array electrodes with the cortical area and RF/nRF locations. **(d).** Example firing rate heatmap of the normalized target-evoked responses to target position A in one marmoset. RF electrodes were defined as having a significant increase in activity relative to a baseline period before target onset. nRF electrodes were defined as the adjacent electrodes that did not show a significant change in activity. **(e).** Same as (d), but for target position B. **(f).** Normalized evoked responses across RF and nRF electrodes. Shaded regions indicate S. E. M.

We first tested whether there was any difference in activity across either the RF or nRF electrodes that correlated with the detection performance of the monkeys. As previously reported, baseline-normalized evoked responses in the RF (100 to 200 msec to target onset) population were significantly stronger on trials when the monkey detected the target (hits) as compared to when the monkey failed to detect the target (hits = 3.22 ± 0.26 S.E.M; misses = 2.81 ± 0.21; p = 0.00034, Wilcoxon signed-rank test; Fig. 3*a, d*). However, we found the opposite effect in the nRF population. nRF activity during the evoked period was significantly greater on misses compared to hits (misses = 1.06 ± 0.041 v. hits = 0.92 ± 0.038 S.E.M.; p = 0.0000050, Wilcoxon signed-rank test; Fig. 3*b, e*).

**Figure 3.**
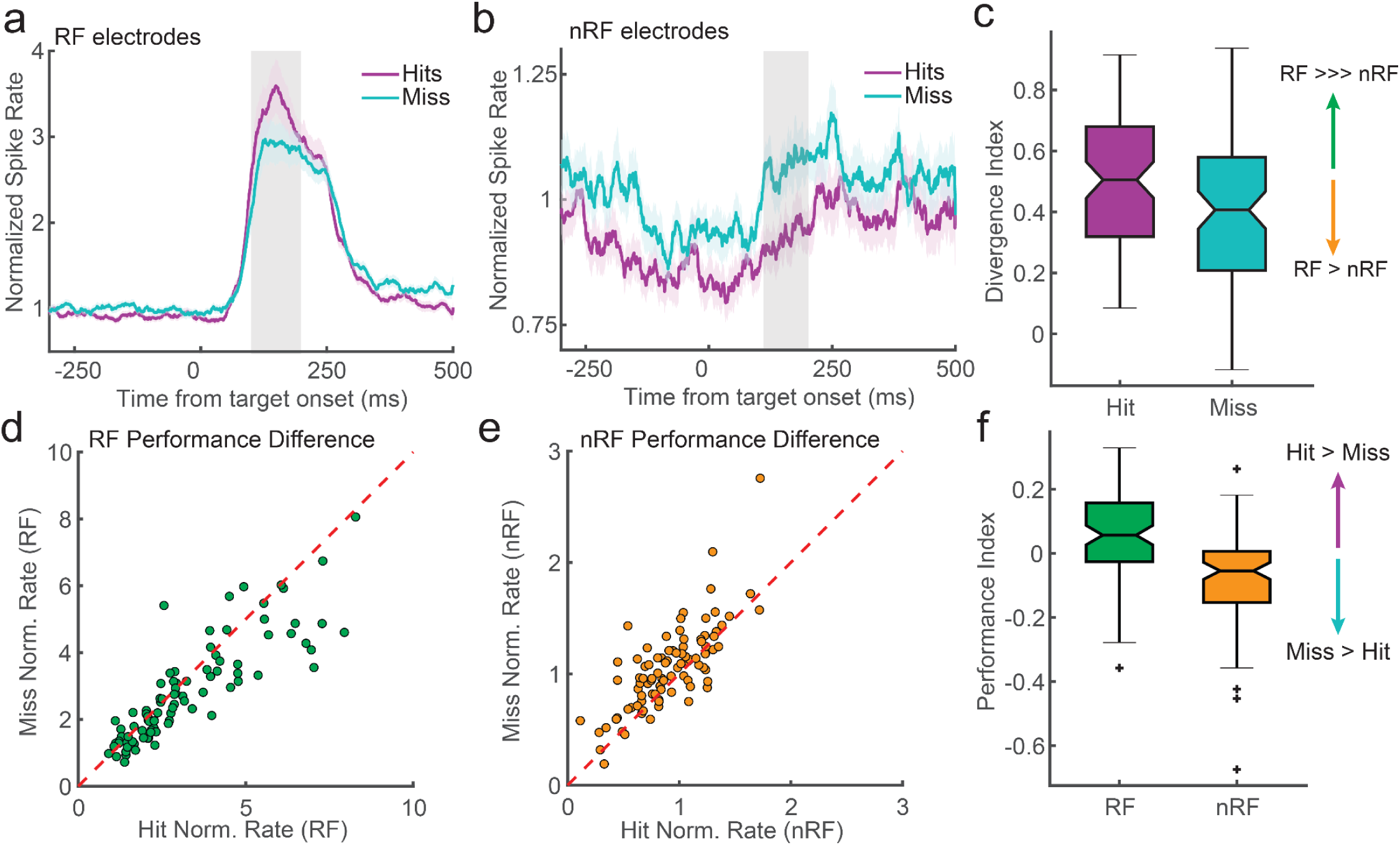
RF and nRF activity divergence is correlated with better detection performance. **(a).** The target-evoked response was stronger across hit trials (purple) compared to miss trials (cyan) for RF electrodes (hits = 3.22 ± 0.26 S.E.M; misses = 2.81 ± 0.21; p = 0.00034, Wilcoxon signed-rank test). Shaded region indicates the window tested (100-200 msec after target onset). **(b).** Intrinsic activity was stronger on miss trials compared to hit trials for nRF electrodes during the same period (misses = 1.06 ± 0.041 v. hits = 0.92 ± 0.038 S.E.M.; p = 0.0000050, Wilcoxon signed-rank test). **(c).** The divergence index (difference over sum) for the RF and nRF populations was significantly greater hit trials compared to miss trials during the evoked response (Hit DI = 0.499 ± 0.02 S.E.M.; Miss DI = 0.390 ± 0.03 S.E.M.; p = 3.5×10^−12^; Wilcoxon rank sum test). **(d).** Scatter plot shows the difference in evoked responses for RF electrodes across recording sessions for hits (x-axis) and misses (y axis). **(e).** Same as (d), but for nRF electrodes. **(f).** The performance index (difference over sum) for hit and miss responses was significantly positive (stronger on hits) for RF electrodes and significantly negative (stronger on misses) across nRF electrodes during the evoked response (RF PI = 0.063 ± 0.01 S.E.M.; p = 0.00051; nRF PI = −0.077 ± 0.01 S.E.M; p = 0.00032; Wilcoxon rank sum test).

We next quantified a Divergence Index (DI; RF-nRF difference over sum) to compare the extent to which RF and nRF activity differed on hit v. miss trials. Values closer to 1/−1 reflect greater divergence and values closer to 0 weaker divergence. Hit trials had significantly stronger divergence between RF and nRF channels than miss trials (hit DI = 0.50 ± 0.023 S.E.M.; miss DI = 0.39 ± 0.26 S.E.M.; p = 3.5×10^−12^, Wilcoxon signed-rank test; Fig. 3*c*). To compare the relative change in activity between RF and nRF populations across hits and misses, we also calculated a Performance Index (PI: hit-miss difference over sum) where positive values reflect stronger activity on hits v. misses, and negative values reflect stronger activity on misses v. hits (Fig. 3*f*). RF PI was significantly greater than zero and nRF PI was significantly less than zero consistent with there being strong RF and weaker nRF activity on hits, and weaker RF and stronger nRF activity on misses (RF PI = 0.063 ± 0.01 S.E.M.; p = 0.00051; nRF PI = −0.077 ± 0.01 S.E.M; p = 0.00032; Wilcoxon rank sum test).

What might give rise to this divergence in population responses between RF and nRF locations on hits and misses? One explanation may be that the stronger evoked responses in the RF population that occurred on hit trials suppressed the activity in the surrounding population via the recruitment of lateral inhibition (Fig. 4*a*), and on miss trials the weaker evoked responses in the RF population recruited less inhibition, permitting more intrinsic activity in the nRF population. If this were true, we would predict there to be a negative correlation between the magnitude of the RF evoked response and the intrinsic activity in the nRF population across hit trials, and this correlation to be weaker across miss trials. However, if it were the case that the divergence between RF and nRF populations were instead driven by shared intrinsic activity states at the time of the target-evoked response (Fig. 4*b*), RF and nRF population responses would move in the same direction, resulting in positive correlations on both hits and misses.

**Figure 4.**
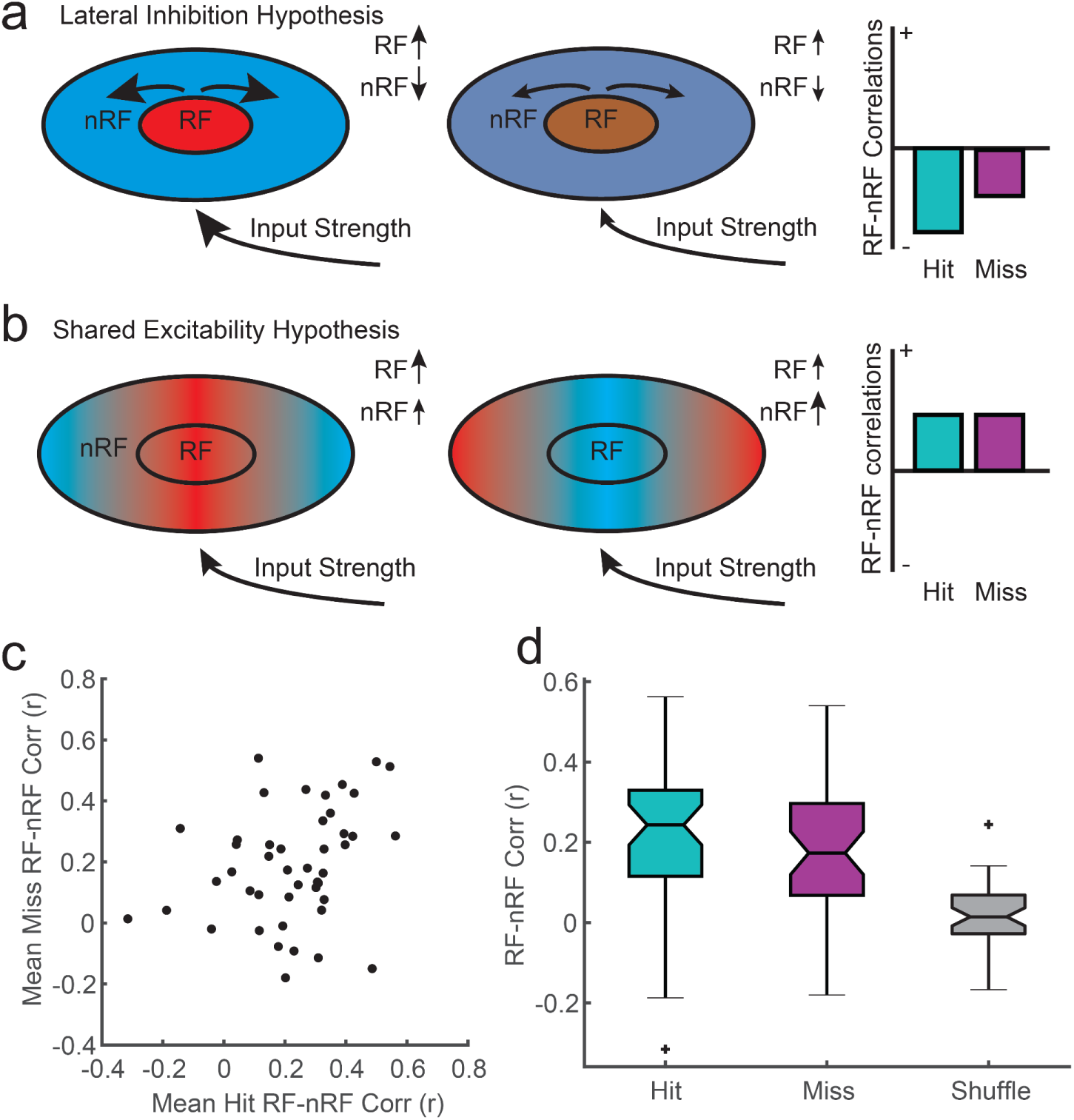
Divergence between RF and nRF activity during the evoked response is not consistent with lateral inhibition. **(a).** Illustration of lateral inhibition hypothesis. Stronger evoked responses will recruit stronger normalizing inhibition, which can suppress the intrinsic activity of the surrounding nRF populations. Weaker evoked responses will recruit less inhibition, resulting in stronger nRF activity. Lateral inhibition would predict negative correlations between RF and nRF activity that are stronger on hits than misses because hits show stronger evoked RF responses. **(b).** The spatiotemporal excitability hypothesis is that the spatial alignment of depolarized states will fall off across the RF and nRF populations, resulting in a degree of shared depolarization state. The consequence of spatially distributed depolarization states are positive correlations that do not differ between hits and misses. **(c).** Distribution of observed correlations between RF and nRF activity across sessions. There was a significant positive correlation across sessions (Pearson’s *r* = 0.46, 95% CI [0.2, 0.66]; p = 0.0012). **(d).** Both hits and misses showed significant positive RF-nRF correlation compared to a shuffle control (p < 0.000001, Wilcoxon signed-rank test). There was no difference in correlation between hits and misses (p = 0.26, Wilcoxon signed-rank test).

To test between these alternatives, we measured the RF-nRF activity correlation across hit and miss trials for each target location across recording sessions (Fig. 4*c*). We found that, across sessions, RF-nRF correlations on hits and misses were significantly positive as compared to the correlations observed after a trial shuffling of RF and nRF activity (mean hit correlation = 0.22 ± 0.03 S.E.M; p = 8.6×10^−10^, Wilcoxon rank sum test; mean miss correlation = 0.18 ± 0.03 S.E.M; p = 1.0×10^−7^, Wilcoxon signed-rank test; Fig. 4*d*). Further, there was no significant difference in the magnitude of correlations between hit and miss trials across sessions (p = 0.26, Wilcoxon signed-rank test). This evidence suggests that the divergence between RF and nRF activity during the evoked response is not due to stronger evoked responses inhibiting the surrounding population.

We next asked if the activity divergence between the RF and nRF populations that correlated with detection performance was temporally specific. To test this, we compared RF and nRF activity in the 200 ms window preceding target onset across hit and miss trials. We found that shared rather than divergent activity states prior to target onset predicted performance. Both RF and nRF populations had significantly higher pre-stimulus activity on miss trials compared to hit trials (RF: misses = 0.99 ± 0.037 vs. hits = 0.91 ± 0.035 S.E.M.; p = 0.00010, Wilcoxon signed-rank test; nRF: misses = 0.96 ± 0.035 vs. hits = 0.87 ± 0.035 S.E.M.; p = 0.0000032, Wilcoxon signed-rank test; Fig. 5*a*–*d*). This suggests that broadly less excitable states prior to target onset facilitate detection performance and divergence between the RF and nRF populations is specific to the evoked response.

**Figure 5.**
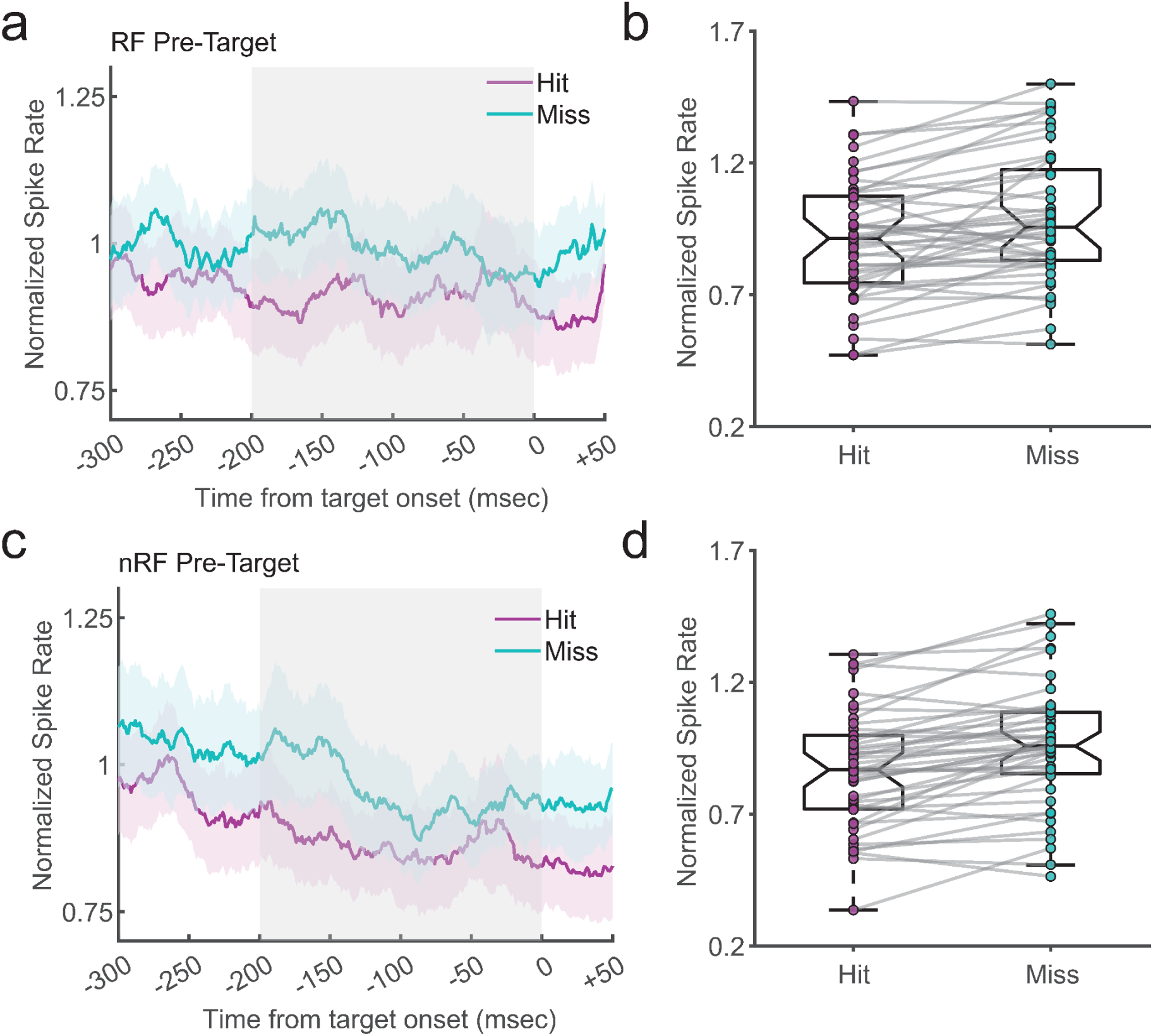
Weak pre-stimulus activity in both RF and nRF populations predicts detection performance. **(a).** RF population activity from −300 to +50 msec with respect to target onset for hit (purple) and miss (cyan) trials. Gray shading indicates the window used for statistical comparison (−200 to 0). **(b)** Boxplots comparing RF activity for hits and misses; misses show significantly greater activity (RF: misses = 0.99 ± 0.037 vs. hits = 0.91 ± 0.035 S.E.M.; p < 0.001, Wilcoxon signed-rank test). **(c).** Same as (a), but for the nRF population. **(d).** Same as (b), but for nRF activity, also showing significantly greater activity on misses (nRF: misses = 0.96 ± 0.035 vs. hits = 0.87 ± 0.035 S.E.M.; p < 0.0001, Wilcoxon signed-rank test).

We next wished to quantify, on a trial-by-trial basis, the relative predictive power of the RF and nRF population activity. We constructed a Generalized Linear Model (GLM) that classified trial outcome (hit, miss) based on the activity of the RF and nRF populations, as well as the RF-nRF divergence both before and after target onset (logit link function, binomial error distribution, N = 4,865 trials). To ensure robust model outcomes we ran the GLM 100 times using randomly shuffled trial assignments for the training and testing set in each iteration. For each run, we computed the area under the curve (AUC) of the receiver operating characteristic (ROC) curve for each model and then averaged the AUC across iterations to assess the predictive power for each variable. We found that in both the pre-stimulus period and the evoked response period, activity in the nRF population was a stronger predictor of trial outcome than the RF activity (mean AUC RF pre = 0.50 ± 0.0004 S.E.M, mean AUC nRF pre = 0.56 ± 0.0003 S.E.M; Cohen’s d = 4.27, mean AUC RF post = 0.53 ± 0.0004 S.E.M, mean AUC nRF post = .57 ± 0.0003 S.E.M; Cohen’s d = 2.33, Fig. 6*a, b*). Further, in the post-stimulus period, the divergence between the RF and nRF activity was more predictive than either the RF or nRF activity in predicting trial outcomes (mean AUC DI post = 0.58 ± 0.001 S.E.M; DI vs. RF Cohen’s d = 2.68, DI vs. nRF Cohen’s d = 0.57; Fig. 6*b, c*). This offers the somewhat surprising result that behavioral performance for detecting faint visual targets from trial to trial depends more on the activity in the neural populations that do not respond to the target rather than on the sensory-evoked response generated by the target-encoding neurons.

**Figure 6.**
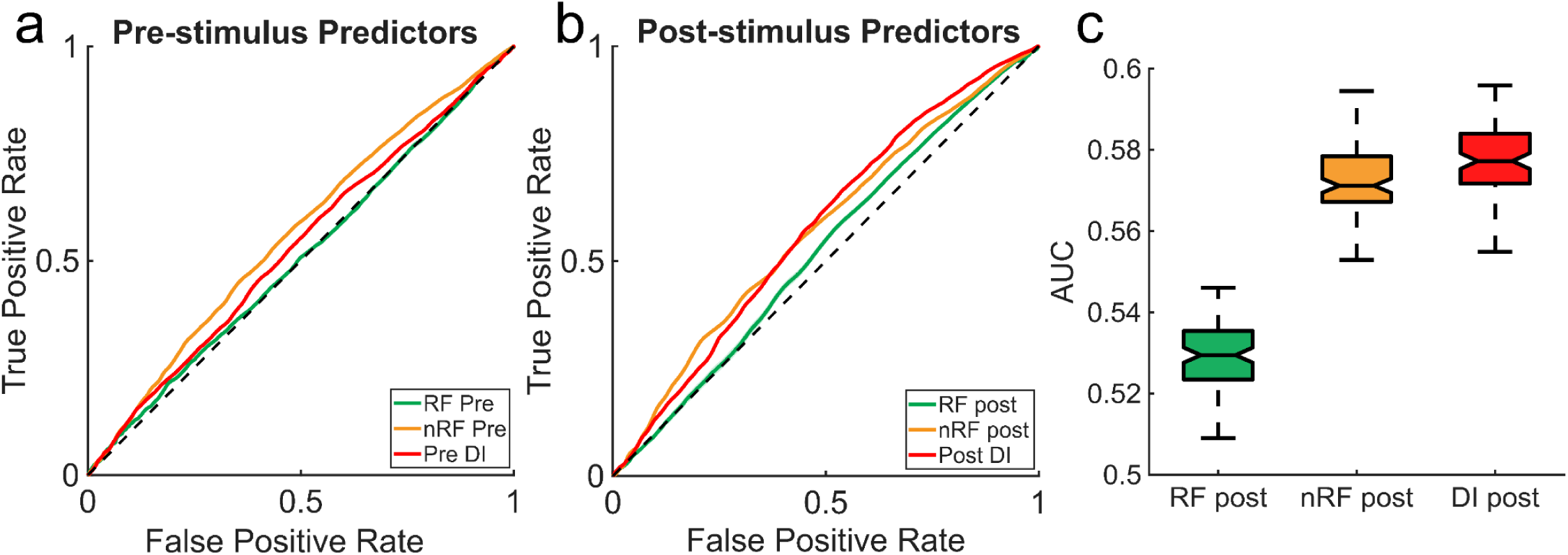
The distributed activity is a stronger predictor of behavioral choice than the RF population activity alone. **(a).** The average ROC curves across 100 GLM interactions using the pre-stimulus RF activity (green line), the pre-stimulus nRF activity (yellow line), and the pre-stimulus divergence index values RF (red line). The black dashed line represents chance performance.. **(b).** The same ROC curves as in a but from the evoked response period. **(c).** Boxplots comparing the distributions of AUC values across the 100 iterations of the GLM trained on activity during the post stimulus period. The divergence shows significantly higher AUC values than the RF and nRF population (RF: 0.53 ± 0.017 S.E.M vs. nRF: 0.57 ± 0.001 S.E.M vs. DI: 0.58 ± 0.001 S.E.M; RF vs. nRF: Cohen’s d = 2.33, RF vs. DI: Cohen’s d = 2.68, nRF vs. DI: Cohen’s d = 0.57).

Finally, we asked whether there was evidence for shared or more heterogeneous states of intrinsic excitability across hits and misses at the time of the target-evoked response. To test this we examined the phase of LFP fluctuations recorded on the RF and nRF electrodes. The LFP reflects changes in the balance of excitatory and inhibitory synaptic activity in the local population around the electrode tip (Okun et al., 2010; Gao et al., 2017). The phase of the LFP can be used as a proxy for changes in intrinsic states of population excitability with π/−π rad phases reflecting more excitable states and 0 rad phases reflecting less excitable states (Mazzoni et al., 2010; Davis et al., 2021). This is validated by the observation that spike probability is often coupled to the phase of broadband LFP fluctuations (Davis et al., 2022). In previous work we found that detection performance depended upon the alignment of intrinsic traveling waves of LFP activity with the RF population, with certain wave states facilitating more excitable states at the time of the evoked response, resulting in better detection performance (Davis et al., 2020). We would therefore predict that there are more excitable LFP phases (e.g. closer to π/−π rad) on nRF channels at the time of the target evoked response on misses compared to hits.

To test this, we calculated the circular mean of the LFP phase filtered from 5-40 Hz across nRF channels. We used a wideband filter on the LFP because spikes during spontaneous activity are coupled to broadband fluctuations in LFP, particularly when no narrowband oscillation is dominant as we find in our recordings from Area MT(Davis et al., 2022). Furthermore, the use of a narrowband filter can distort estimates of signal phase when the frequency content of the LFP falls outside the filter band. The lower and upper bounds selected were chosen to reduce the contribution of large amplitude low frequency components often associated with arousal and higher frequency components that can contain spike artifacts that induce spurious spike-LFP phase coupling (Zanos et al., 2011; Waldert et al., 2013; Ray, 2015).

We found that, at the start of the target-evoked response (~70 msec) the mean phase of the LFP over nRF electrodes significantly diverged between hits and misses with misses showing phases closer to π rad consistent with the nRF population being in a more excitable state on misses (mean hit phase = 1.27 ± 0.11 rad v. mean miss phase = 2.85 ± 0.12 rad; p < 0.00001 Watson-Williams test). Fig. 7). RF activity was not included in this analysis as the distribution of phases was uniform at this time, meaning there was no significant mean phase (p = 0.51; Rayleigh test for non-uniformity). These results are consistent with our observation that the intrinsic excitability of the nRF population, specifically at the time of the evoked response, shapes the separation between RF and nRF population activity and modulates the likelihood the monkey will detect the target.

**Figure 7.**
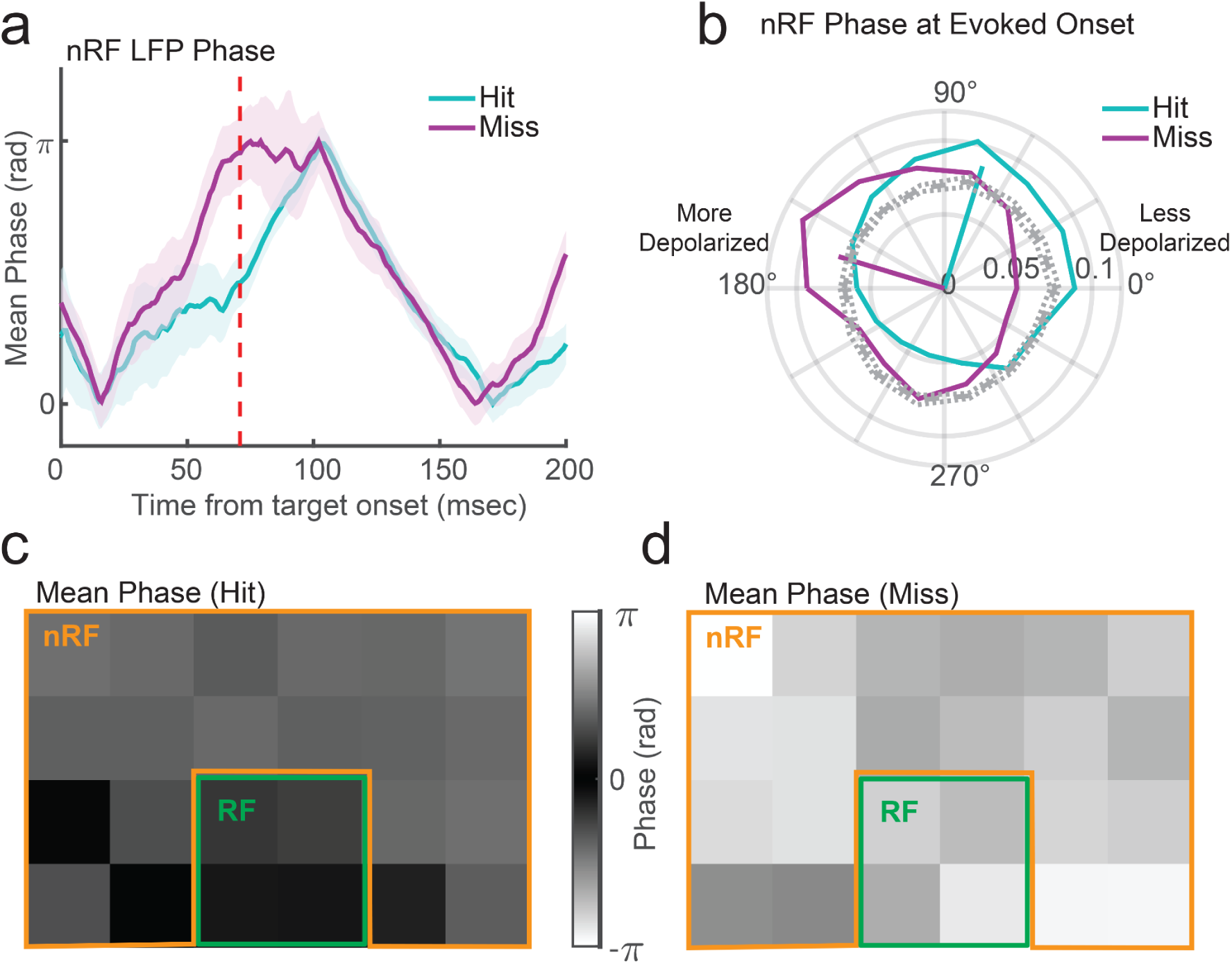
nRF depolarization state, as indexed by LFP phase, predicts performance at the onset of the evoked response. **(a).** The average LFP phase (abs. value) on nRF channels is closer to pi (e.g. more excitable state) on miss trials v. hit trials at the onset of the evoked response (red dashed line). **(b).** nRF electrodes were significantly more depolarized (e.g. closer to π/−π rad phase) on misses compared to hits at 60-70 msec after target onset (mean hit phase = 1.27 rad v. mean miss phase = 2.85 rad; p < 0.00001 Watson-Williams test). Data not shown for RF channels as there was no significant phase organization across misses (p = 0.51; Rayleigh test for non-uniformity). **(c-d).** The mean phase map on hit and miss trials for RF and nRF electrodes collapsed across target locations and monkeys. The nRF channels are significantly darker (less depolarized) on hits compared to misses just prior to the onset of the evoked response.

## Discussion

In this work we wished to understand how variable states of intrinsic cortical activity contribute to detection thresholds in the visual system. It is well understood that stronger sensory-evoked responses are better detected. However, what determines whether a sensory response is strong enough to be detected? If the threshold for detection is drawn from the expected activity of the cortical population, then moments when cortical activity is relatively silent should facilitate detection by reducing the response criterion (Parker and Newsome, 1998; Iemi et al., 2017). However, if these quiescent states reduce the excitability of sensory-responsive neurons, then that would reduce their evoked response. Therefore, the contribution of intrinsic activity on detection thresholds depends on the degree to which the intrinsic activity is shared or divergent across the populations that form the response criterion and represent the sensory signal.

Variability in cortical activity can largely be attributed to the variable states of the numerous synaptic inputs that neurons receive (Steriade et al., 2001; Renart et al., 2010). It has been argued that shared sources of variability common across the population (e.g. states of arousal, locomotion, etc) are orthogonal to sensory representations in cortical populations and might have negligible impact on decoder performance (Stringer et al., 2019, 2021; Rumyantsev et al., 2020). However, animal behavior often underperforms the fidelity of the sensory representation encoded in the population (Beaman et al., 2017; Panzeri et al., 2017; Stringer et al., 2021; Russell et al., 2024). Why are choice probability signals often so weak? There is evidence that behavior does not rely solely on the sensory representation in RF populations, but is sampling across heterogeneous populations spanning varying sensory dimensions (Bounds and Adesnik, 2025). The majority of the inputs to neurons come from recurrent connections within cortical networks that are shared across dimensions of space and feature shaping correlations based on proximity and tuning similarity across the cortical area (Bosking et al., 1997; Smith and Kohn, 2008; Markov et al., 2011). Variable activity amongst these connections forms heterogeneous spatiotemporal fluctuations that can impact detection performance depending on how they align with encoding and non-encoding populations (Davis et al., 2020, 2024).

We propose a framework wherein spatiotemporally heterogeneous states of intrinsic excitability across a cortical area can drive stronger evoked responses in stimulus responsive populations (i.e. RF populations) while at the same time reducing the level of activity in the surrounding, nRF population. When such a state occurs, there is greater divergence between the RF and nRF activity that facilitates the detection of the stimulus from the background activity in a downstream integrator. We find evidence consistent with that framework. The magnitude of the RF population target-evoked response is positively correlated with target detection and the magnitude of spontaneous activity in the nRF population is negatively correlated with target detection. The divergence between RF and nRF population activity correlates with the trial-to-trial performance of the monkey. This divergence is specific to the period of the evoked response. Prior to the response, a general state of quiescence better predicts detection performance.

Our results are consistent with prior work from the field. Fluctuations in spiking activity and local field potentials recorded intracranially and scalp field potentials measured with EEG or BOLD signals measured with fMRI both show that the pre-stimulus state of activity is predictive of detection performance (Hesselmann et al., 2008; Wu et al., 2024). States that facilitate stronger evoked potentials correlate with better sensitivity, and states that reflect weaker intrinsic activity correlate with shifts in the threshold for reporting detection. However, EEG and fMRI measures lack the spatial resolution to segregate RF and nRF population activity during either the pre-stimulus or stimulus-evoked period. We find that comparing the state of activity across these two populations is more predictive than looking at the evoked response magnitude alone.

Our framework offers a mechanistic interpretation for why states of low arousal may impair detection sensitivity. During states of low arousal, cortical activity is more correlated in space and time, exhibiting increases in large-amplitude and low-frequency shared fluctuations across the cortex (McGinley et al., 2015). These shared fluctuations would reduce the divergence between RF and nRF populations, resulting in a need for stronger evoked responses to breach the threshold for detection. Alternatively, when arousal is higher as occurs with states of attentiveness, the reduction in low frequency correlated variability across the population would decouple RF and nRF population activity, permitting divergence that facilitates detection sensitivity (Cohen and Maunsell, 2009; Mitchell et al., 2009; Schölvinck et al., 2015). We speculate that one of the benefits of attention may be to align excitability topologies across a cortical area that drives the divergence of RF and nRF populations depending on the task at hand.

We considered two possible mechanisms that might drive the difference in RF-nRF divergence across hits and misses in our task. The first mechanism was lateral inhibition. A stronger stimulus-evoked response should drive stronger normalizing inhibition, which could suppress the surrounding nRF population and create an activity divergence that facilitates stimulus detection. However, if this were the explanation for our observations, we would expect to find a negative correlation between RF and nRF responses across trials. Instead we found a positive one. This suggests the greater divergence we see on hits can not be due to suppression caused by the stimulus, but rather due to interactions between intrinsic and sensory-evoked activity.

The second mechanism we tested was the presence of a less excited state of intrinsic activity in the time leading up to the target-evoked response. We found that LFP phases, an index of relative population excitability, was correlated with detection performance in the nRF population at the time of the target response. In addition we found that weaker nRF activity before target onset correlated with stronger divergence post-stimulus. Here we show that these effects are spatially specific, rather than homogeneous across the cortical population.

In summary, we find that the state of intrinsic activity plays a role in contextualizing sensory responses during sensory detection. While the sensory information encoded by neuronal populations is necessary for detection computations, the behavior of the animal is better predicted by the intrinsic activity of the larger population. This is consistent with detection computations comparing the intrinsic and evoked activity with stronger divergence between RF and nRF populations facilitating better detection sensitivity. Our results support the view that, while sensory signals are best encoded by the receptive population, behavior is better predicted by ensembles that span stimulus encoding and non-encoding populations.

## Materials and Methods

The data analyzed in this work was collected from two adult marmoset monkeys (*Callithrix jacchus*; one male: monkey W and one female: monkey T). All surgical procedures were performed with the monkeys under general anesthesia in an aseptic environment. All experimental methods were approved by the Institutional Animal Care and Use Committee (IACUC) of the Salk Institute for Biological Studies and conformed with NIH guidelines (protocol 14-00014).

### Electrophysiological Recordings

Each monkey was surgically implanted with a headpost for head stabilization and eye tracking. A craniotomy and duratomy was made over Area MT (stereotaxic coordinates 2 mm anterior, 12 mm dorsal). An 8×8 (64 channel, 400 μm spacing with a pitch depth of 1.5 mm; monkey W) and 9×9 with alternating channels removed (40 channel, monkey T) Utah array was implanted using a pneumatic inserter wand (Blackrock Microsystems, UT). The craniotomy was closed with Duraseal (Integra Life Sciences, monkey W) or Duragen (Integra Life Sciences, monkey T), and covered with a titanium mesh embedded in dental acrylic.

Multielectrode recordings from the Utah arrays were made via 2 32-channel Intan RHD 2132 headstages connected to an Intan RHD2000 USB interface board (Intan Technologies LLC, Los Angeles, USA) controlled by a Windows computer. Data was sampled at 30 kHz from all channels. Digital and analog signals were coordinated through National Instrument DAQ cards (NI PCI6621) and BNC breakout boxes (NI BNC2090A). Neural data was broken into two streams for offline processing of spikes (single-unit and multi-unit activity) and LFPs. Spike data was high-pass filtered at 500 Hz and candidate spike waveforms were defined as exceeding 4 standard deviations of a sliding 1 second window of ongoing voltage fluctuations. Artifacts were rejected if appearing synchronously (within 0.5 ms) on over a quarter of all recorded channels. Segments of data (1.5 ms) around the time of candidate spikes were selected for spike sorting using principal component analysis through the open source spike sorting software MClust in Matlab (A. David Redish, University of Minnesota). Sorted units were classified as single- or multi-units, or noise and single units were validated by the presence of a clear refractory period in the autocorrelogram. Units classified as noise were eliminated from analysis. Spiking activity from single and multi-unit spikes recorded on the same channel were combined to give a single channel spike count. LFP data was low-pass filtered at 300 Hz and down-sampled to 1000 Hz.

### Experimental Design and Statistical Analysis

#### Detection Task

Marmosets were trained to enter a custom-built marmoset chair that was placed inside a faraday box with an LCD monitor (ASUS VG248QE) at a distance of 40 cm. The monitor was set to a refresh rate of 100 Hz and gamma corrected with a mean gray luminance of 75 candelas/m^2^. The marmosets were headfixed by the headpost for all recordings. Eye position was measured with an IScan CCD infrared camera sampling at 500 Hz. The MonkeyLogic software package developed in MATLAB (https://monkeylogic.nimh.nih.gov/)(Asaad et al., 2013) was used for stimulus presentation, behavioral control, and recording of eye position.

The marmosets were trained to saccade to a marmoset face to initiate each trial. Upon the gaze arriving at the face, it disappeared and was replaced with a white fixation point (0.15 DVA). The marmosets held fixation on the fixation point (1.5 visual degree tolerance) for a minimum delay (400 ms monkey W, 300 ms monkey T) awaiting the appearance of a drifting Gabor target (4 DVA diameter; appearing 6-7 DVA eccentricity at 1 of 2 equally eccentric locations in the visual field contralateral to the recording array). The target contrast was titrated to a value that each monkey detected approximately 50 percent of the time (1-2% Michelson contrast). The onset time of the target (after the minimum delay) was drawn from an exponential distribution (mean 200 msec) to yield a flat hazard function. After the delay, the target appeared for 200 msec and the monkey had 500 msec from target onset to saccade to the location of the target for a juice reward. Early fixation breaks (defined by the excursion of the eye position from the fixation window) were excluded from analysis. High contrast probes (10% Michelson contrast) randomly occurred on 10 percent of trials. If the monkey’s performance on these easy target trials was below 70% within a session that session was excluded from analysis. We analyzed a total of 18 sessions from monkey T and 27 sessions from monkey W for a total of 45 sessions.

To ensure marmosets did not adopt a guessing strategy, 10 percent of trials were catch trials where no target appeared and marmosets were rewarded for maintaining fixation through the duration of the trial. Performance on these catch trials was above 70 percent for both monkeys (monkey W: 71% correct reject; monkey T: 73% correct reject). Unlike a 2-alternative forced choice task, where chance performance is 50 percent, because of the unpredictable onset time of the target and the presence of catch trials, we estimate chance performance on this task to be approximately 22 percent based on Monte Carlo simulations with random guessing strategies. Further, the reaction time distributions for both monkeys were normal with respect to target onset, and their false alarm rates were low suggesting they were faithfully performing the task as intended.

#### Receptive field mapping

MT receptive fields were mapped by calculating the spike-triggered average from the reverse correlation of visual probes presented on the computer monitor. The marmoset began a mapping trial by fixating an image of a marmoset face. After holding fixation for 200 ms, a drifting Gabor target (50 percent Michelson contrast; spatial frequency: 0.5 cycles per degree; temporal frequency: 10 cycles per second; drifting in one of 8 (monkey W) or 6 (monkey T) possible directions) appeared at a random location on the screen contralateral to the implanted array. After 200 ms the Gabor disappeared and, after a random delay drawn from an exponential distribution (mean delay 30 ms) reappeared at another random location. The processes repeated until the monkey broke fixation (defined as an excursion from the fixation zone beyond 1.5 d.v.a.). The monkey received a juice reward proportional to the number of probes presented on the receptive field mapping trial. We calculated both the spatial location of the receptive field of single and multi-unit activity on the Utah array as well as the tuning curves for the preference for motion direction of the probes. Based on receptive field locations, the top 32 channels of Monkey W’s Utah array were excluded from analysis as these channels were likely to be outside of Area MT.

#### RF and nRF definition

RF and nRF channels were defined based on significant increases in firing rate to the presentation of the target at one of the two target locations. The spike rate during the target-evoked response (100-200 msec after target onset) was compared to the average baseline spike rate over the interval 300 msec prior to the appearance of the target. Channels with a significant increase in firing rate across sessions were defined as RF channels for that location. In both monkeys there were 2 RF channels at each target location. nRF channels were defined by finding channels 1 location away from RF channels that did not have a significant change in firing rate during the evoked period. We excluded channels that had baseline firing rates below 1 Hz. For monkey T there were 6 nRF channels for each target location, and for monkey W there were 5 nRF channels for the lower target location and 6 nRF channels for the upper target location. We also considered more distant channels as an alternative nRF population (farRF) which were an additional 1 channel more distant from the nRF population, as well as the alternative target location, which consisted of the RF channels at the location where the target did not occur.

#### Divergence Index

We calculated an index that captured the divergence between RF and nRF activity across recording sessions. The average spiking activity on each channel was baseline normalized to the 300 msec prior to target onset. The normalized evoked response (100-200 msec after target onset) was averaged across RF and nRF channels respectively. DI was calculated separately for hit miss trials as:

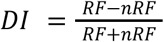

A similar Performance Index was calculated by comparing across hit and miss trials within RF and nRF channels.

#### Local Field Potential Analysis

Local field potential (LFP) data were analyzed from the period 300 msec before target onset to the period 300 msec after target onset. The raw LFP was filtered from 5-40 Hz (4th order Butterworth forward-reverse filter using the *filtfilt* function in MATLAB). LFP power < 5 Hz was filtered out to exclude low frequency fluctuations that are associated with slow changes such as changes in arousal(Steriade et al., 2001; Petersen et al., 2003; McGinley et al., 2015). LFP power > 40 Hz was excluded to avoid the possibility of bleedthrough of spiking into the LFP, which could potentially lead to spurious spike-LFP coupling(Zanos et al., 2011). We then calculated the *Generalized Phase* (GP) of this wideband signal using previously described methods for calculating phase of signals with relatively broad spectral content. GP represents an advancement over the standard Hilbert Transform as it corrects for the breakdowns that can occur when calculating the instantaneous phase of a signal with broad spectral content.

Peri-target phase maps were taken at the time of the average target-evoked onset (70 msec after the target appeared). First, we took the phase distribution on each channel and tested whether the distribution was uniform (Rayleigh’s test for non-uniformity). RF channels did not show a significant mean phase direction and were therefore not considered further. nRF channels did show a phase preference. We calculated the circular difference between the mean phase on nRF channels across hit and miss trials and π radians.

#### Generalized Linear Model

We fit a generalized linear model (GLM) to the average baseline normalized firing rate before (−200 msec to target onset) and during the evoked response (100-200 msec) on each trial across RF and nRF channels. Trials where there were no spikes on either the RF or nRF populations during the response window were excluded from the analysis. The GLM used a binomial model with a logit link function to predict whether each trial was a hit or a miss from the regression of the channel firing rates. Model weights were measured using a random half of the dataset of trials across all recording sessions (matched 50/50 hits and misses) and the model was evaluated by testing the area under the curve (AUC) of the receiver-operator characteristic (ROC) from the left-out half of the data. This process was repeated 100 times to generate a distribution of model performance using either RF or nRF channels, or their combination as separate variables.

## QUANTIFICATION AND STATISTICAL ANALYSIS

Quantification and statistical analysis was performed using MATLAB statistical functions and code from the MATLAB Circular Statistics Toolbox (P. Berens, CircStat: A Matlab Toolbox for Circular Statistics, Journal of Statistical Software, Volume 31, Issue 10, 2009). Details on the Ns, measurements of center and dispersion, generation of null hypotheses, statistical test used, and measurement of significance for each analysis can be found where those analyses are described in the corresponding results and method detail sections of the manuscript.

## Data and code availability

- All data used in this paper will be shared by the lead contact upon request.
- Any additional information required to analyze the data reported in this paper is available from the lead contact upon request.

## Acknowledgements

This work was supported by National Institutes of Health Core Grant (EY014800), the Whitehall Foundation, an Unrestricted Grant from Research to Prevent Blindness, New York, NY, to the Department of Ophthalmology & Visual Sciences, University of Utah, and a National Institutes of Health Institutional NRSA T32 Training Grant T32 EY024234 (Vision Research Training Program, University of Utah).

## Author Contributions

Designed research: Z.W.D.; Performed research: Z.W.D.; Analyzed data: Z.W.D., D.J.; Wrote the paper: Z.W.D., D.J.

## Conflicts of Interest

The authors declare no conflicts of interest.

## Notes

### Competing Interest Statement

The authors have declared no competing interest.

